# Non-Markovian memory in a bacterium

**DOI:** 10.1101/2023.05.27.542601

**Authors:** Kunaal Joshi, Karl F Ziegler, Shaswata Roy, Charles S Wright, Rhea Gandhi, Jack Stonecipher, Rudro R Biswas, Srividya Iyer-Biswas

## Abstract

Do individual bacterial cells retain memories of the history of environmental conditions experienced in previous generations? Here we directly address this question through a synthesis of physics theory and high-precision experiments on statistically identical, non-interacting individual bacterial cells, which grow and divide with intrinsic stochasticity in precisely controlled conditions. From these data, we extract “emergent simplicities” in the seemingly complex interplay between history dependence, persistence, and transience in the stochastic memories of the dynamic environments experienced by individuals over multiple generations. First, we find that the instantaneous single-cell growth rate is the key physiologically relevant quantity where intergenerational memory is stored. In contrast, the cell size dynamics are memory free, or Markovian, over intergenerational timescales. Next, we find that the effect of experiencing dynamic environments can be captured quantitatively by recalibrating the cellular unit of time by the measured mean instantaneous growth rate; the dynamically rescaled cell age distributions undergo a scaling collapse. Moreover, in a given condition, an individual bacterial cell retains history-dependent, or non-Markovian, memory of its growth rate over tens of generations. We derive from first principles a physically-motivated metric to quantify the degree of non-Markovianity. Furthermore, when conditions change, the instantaneous single-cell growth distribution becomes bimodal, as the bacterium’s memory of past environment encountered is reset stochastically and plastically, prior to achieving a new homeostasis.

## Introduction

Homeostasis—the constancy of an organism’s “internal milieu” —is so pervasive a notion in biology and medicine that it has effectively served as an organizational principle for over a century (1–8). Yet, the organism’s “stable” internal environment must also accommodate the need for dynamic physiological adaption under changing external conditions (9–12). How are these seemingly opposing demands to be reconciled by an organism?

Sterling and coworkers have argued that the requirement for organismal homeostasis is conceptually limiting, even flawed (9, 10, 13). Since an organism’s priority is to maximize the possibilities for survival and reproduction, they argue that responsiveness to dynamic external conditions is the primary focus of organismal control, rather than a rigid commitment to pre-determined “set-point” values of homeostatic physiological quantities. Sterling thus advocates for replacing the notion of homeostasis by that of allostasis, wherein the organism achieves “stability through change” by using prior information to learn, anticipate and respond strategically to impending changes. These changes are computed based on historic information, overriding as needed the local error-correction-through-negative feedback routines, which constitute the the presumed mechanism for sustaining homeostasis (9).

Yet bacteria, unicellular organisms without dedicated nervous systems, manage to thrive in changing environmental conditions. How do they reconcile the need to maintain internal stability with plastic responsiveness? What principles govern how a bacterium responds to an exposure to a novel environment? Here, we tackle these questions directly through high-precision experiments on individual bacterial cells experiencing identical, precisely controlled dynamic growth conditions. We characterize the long-term multigenerational stochastic growth and division dynamics of individual *Caulobacter crescentus* cells in both constant and time-varying conditions using our SChemostat technology (Fig. 1) (14–17). At each (asymmetric) division the immobilized stalked cell is retained, while the motile swarmer cell is flowed away in a precisely controlled microfluidic environment. Thus, crowding of the region of interest and physical contact between individuals, unavoidable in other single-cell technologies, is naturally obviated in the SChemostat (14, 18–21). Individual cells are non-interacting, statistically identical and isogenic, and experience identical environmental conditions at a given moment in time, while the conditions themselves can be precisely changed as functions of time using programmable and automated microfluidic controls. Rapid phase-contrast imaging of individual cells at 50– 100 times per cell cycle, in conjunction with the fact that each individual cell can be readily tracked for tens of generations, allows for precise characterization of each individual cell’s size growth rate over a large dynamic range, spanning intra- and inter-generational timescales (for instance, as shown in Figs 1, 2, and 6).

**Fig. 1.**
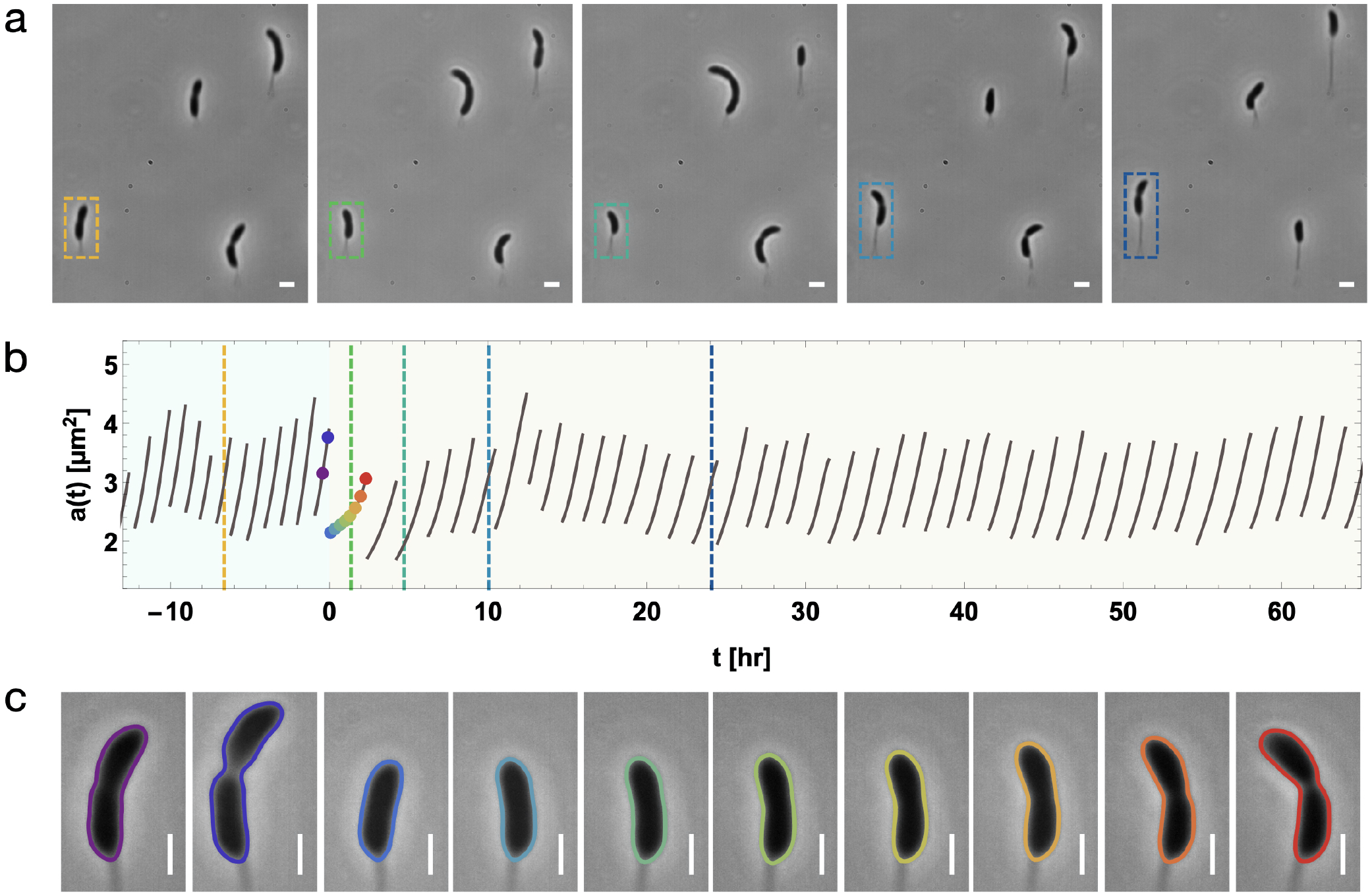
High-precision imaging of multigenerational stochastic growth and division dynamics of individual cells in time-varying conditions using the SChemostat technology. **(a)** Snapshots of a typical field of view at five different time points in the SChemostat setup, containing four cells where the dashed box highlights the same cell with colors corresponding to the times indicated by dashed lines on the trajectory in (b). **(b)** The time evolution of the experimentally measured area growth trajectory of the highlighted cell in (a), as the growth media is abruptly switched from complex (light blue background) to minimal (light yellow background), with the switch occuring at *t* = 0. **(c)** Zoomed-in images of the highlighted cell in (b), superimposed with splines calculated by automated analysis, at approximately equally spaced time points chosen from generations immediately preceding and succeeding the switch in growth conditions. These times are indicated by points with corresponding colors in (b). Using these images and the equal time-spacing between images within each generation, the significant slowing down of cell growth immediately after the switch is visually evident.

**Fig. 2.**
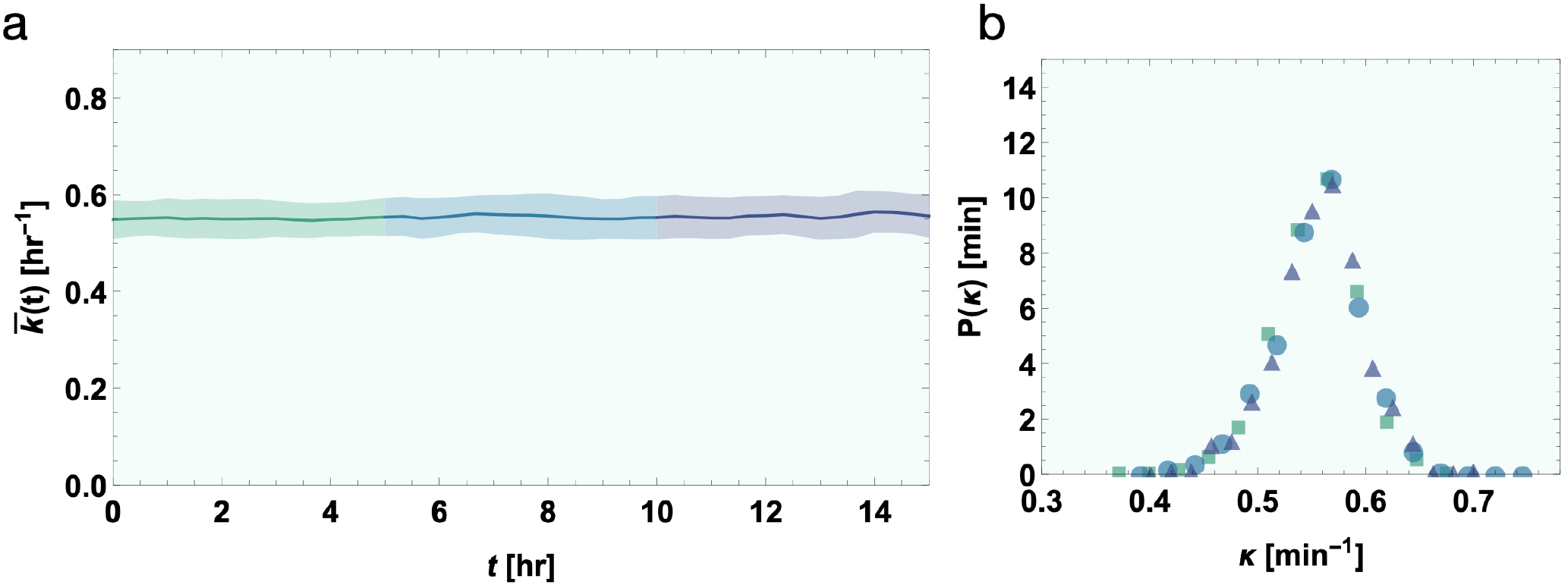
Stochastic intergenerational homeostasis of the individual cell (size) growth rate in constant growth conditions. **(a)** The mean and standard deviation (SD) of asynchronous growth rate are plotted as a function of experiment time in complex media; these are time-variant. **(b)** Distribution of single-cell growth rates plotted for the three different epochs of experimental time (different distributions are color-coded to match the experimental time epochs shown in (a)). These distributions are identical for each epoch, implying the growth rate is in homeostasis.

Our high-precision data reveal that an individual cell’s specific (size) growth rate serves as a repository of intergenerational memory: the dynamics are history-dependent, or non-Markovian (Fig. 3). This stands in contrast to the history-independent, or Markovian, dynamics of size at birth, wherein the initial size of the next generation is solely determined by the current generation’s initial size, and is indepen-dent of the initial sizes of past generations (22); the growth rate of the next generation is affected both by the growth rate of the current generation, as well as that of the preceding generations. We introduce a first-principles-based metric of non-Markovianity (Eq. Eq. (1)) and establish that a cell probabilistically retains memory of its growth rate for tens of generations under constant conditions (Fig. 4). Distinct biphasic signatures of short- and long-term memory characterize the persistence curve (Fig. 5 d–f). In dynamic environments, the mean instantaneous cell size growth rate—surprisingly identical for cell subpopulations experiencing environmental changes at differing phases of their cell cycles—provides a direct calibration of the dynamic cellular unit of time (Fig. 6). Rescaling in turn by this value results in a scaling collapse of the time-dependent cell-age distributions, in remarkable agreement with predictions from theory obtained with no fitting parameters (Fig. 7). Remarkable changes are observed in the distributions of individual cell growth rates, however, which are otherwise in stochastic intergenerational homeostasis in steady state (Fig. 2). When external conditions change, the transient growth-rate distributions become bimodal as the cell’s memory is partially reset, en route to attaining a new state of homeostasis (Fig. 8 a, b). Thus, the cell’s intergenerational memory encoded in its growth rate is reset upon experiencing dynamical changes in environmental conditions (Fig. 10).

**Fig. 3.**
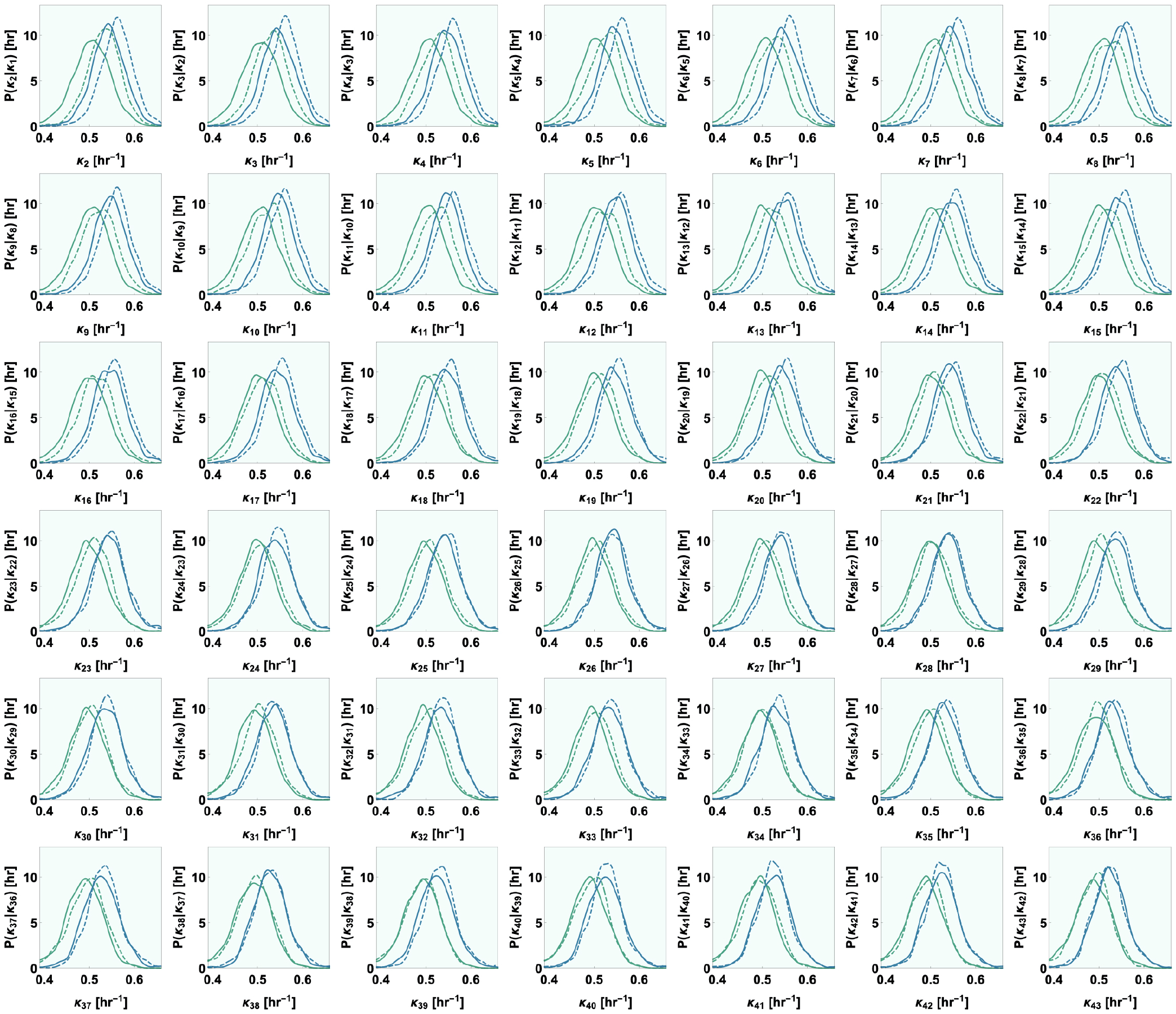
Non-Markovian memory retention for tens of generations in the individual cell (size) growth rate. For cells growing in complex media in steady state, the conditional distributions of the n^th^ generation’s growth rates (*k*_*n*_) are plotted, conditional on whether the previous generation’s growth rate (*k*_*n−*1_) is greater than (blue) or less than (green) the median growth rate, and whether the initial generation’s growth rate (*k*_0_) is greater than (dashed) or less than (solid) the median growth rate. For a completely Markovian process, the distribution of *k*_*n*_ conditional on *k*_*n−*1_ would be independent of the growth rates of all generations prior to *n −* 1, including *k*_0_. Thus, the difference between solid and dashed curves indicates the presence of inter-generational memory in growth rate. These plots show the distributions for *n* ranging from 2 to 43, and it is clear that memory is maintained for over 20 generations.

**Fig. 4.**
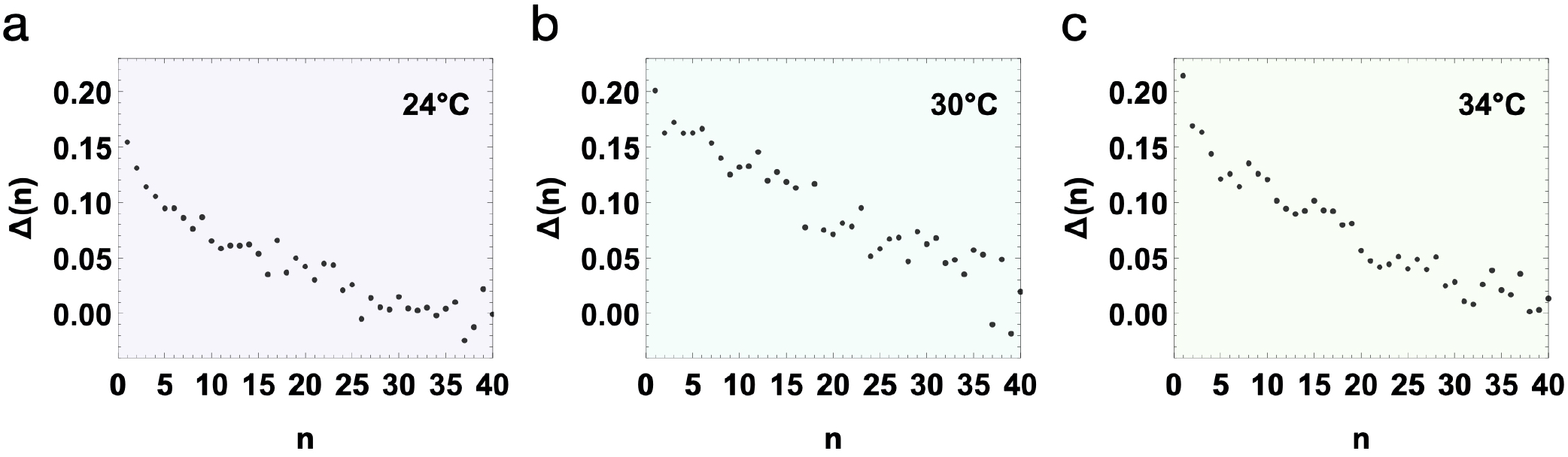
Quantifying the degree of non-Markovianity in the intergenerational single-cell growth-rate dynamics; memory is retained for *∼*40 generations. Δ(*n*), a measure of the non-Markovian behavior of single cell growth rate given by Eq. Eq. (1), is plotted as a function of generation *n*, for cells in steady state at **(a)** 24^*°*^C, **(b)** 30^*°*^C, and **(c)** 34^*°*^C. Δ(*n*) is zero for all *n* for Markovian systems; a non-zero value for Δ(*n*) indicates the presence of non-Markovian memory up to generation *n*. Thus, memory persists until *∼* 40 generations. The positive values of Δ indicate a positive relation between growth rate in the current generation with growth rates of past generations upon compensating for the effect of growth rate in the previous generation.

**Fig. 5.**
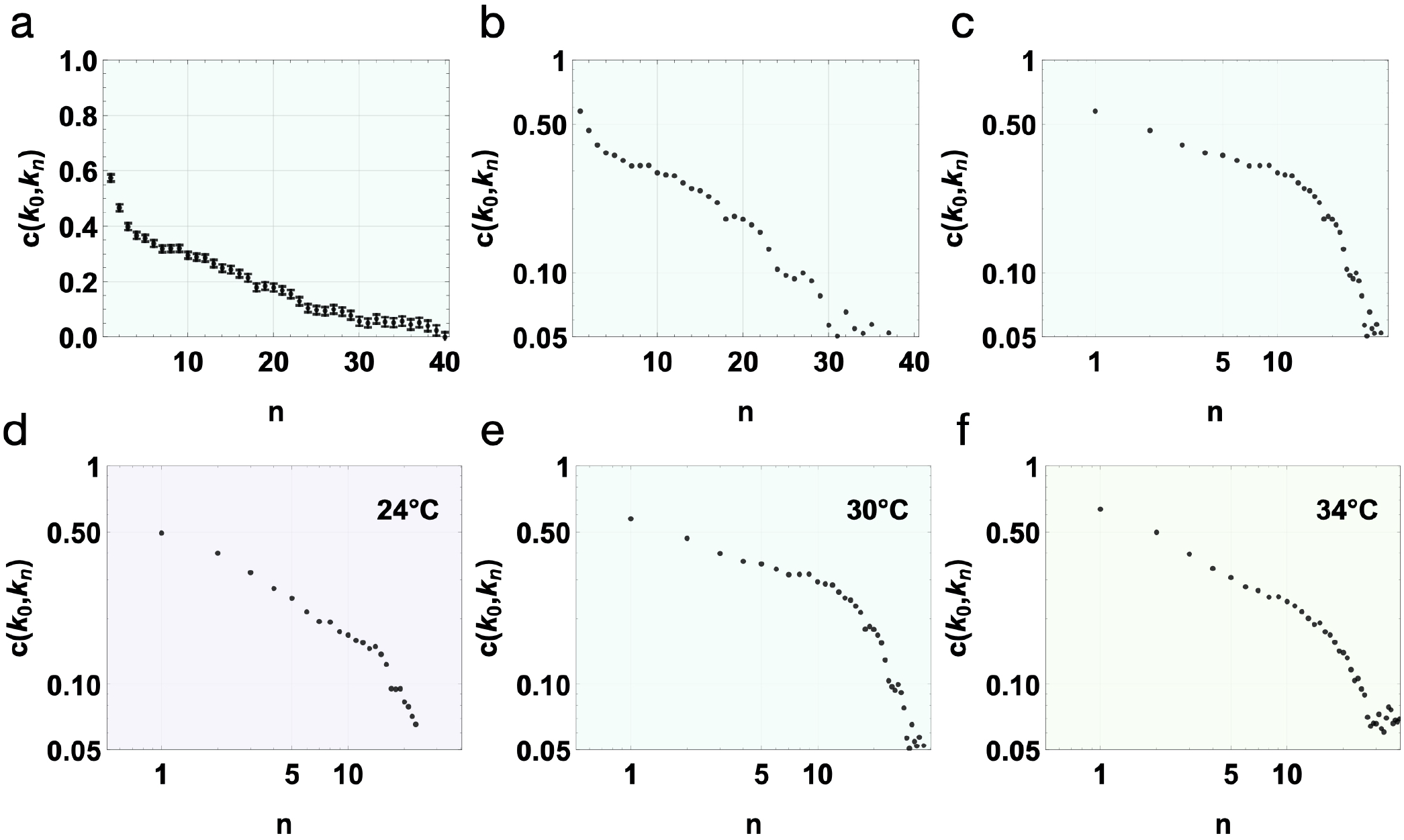
The memory persistence curve extends over tens of generations and displays distinct signatures of long- and short-term memory. **(a–c)** For cells in steady state in complex media at 30^*°*^C, the coefficient of correlation between the growth rate of initial (zeroth) generation and n^th^ generation is plotted as a function of *n*, on a (a) linear scale, (b) log-linear scale, and (c) log-log scale. Approximately 30 generations are required for the correlations to die out, and thus for the cells to forget their initial growth rates. The observation that the trend for small *n* is not a straight line on the log-linear scale indicates that the inter-generational growth rate is non-Markovian. The trend on the log-log scale is a straight line for low *n*, indicating power law behavior. **(d–f)** For cells in steady state in complex media at three different temperatures, the same coefficient of correlation is plotted on a log-log scale. We observe a straight line for low *n* for all three temperatures, indicating that the power law behavior is ubiquitous. The power law changes abruptly across *∼* 20 generations, delineating two distinct regimes corresponding to short-term and long-term memory retention.

**Fig. 6.**
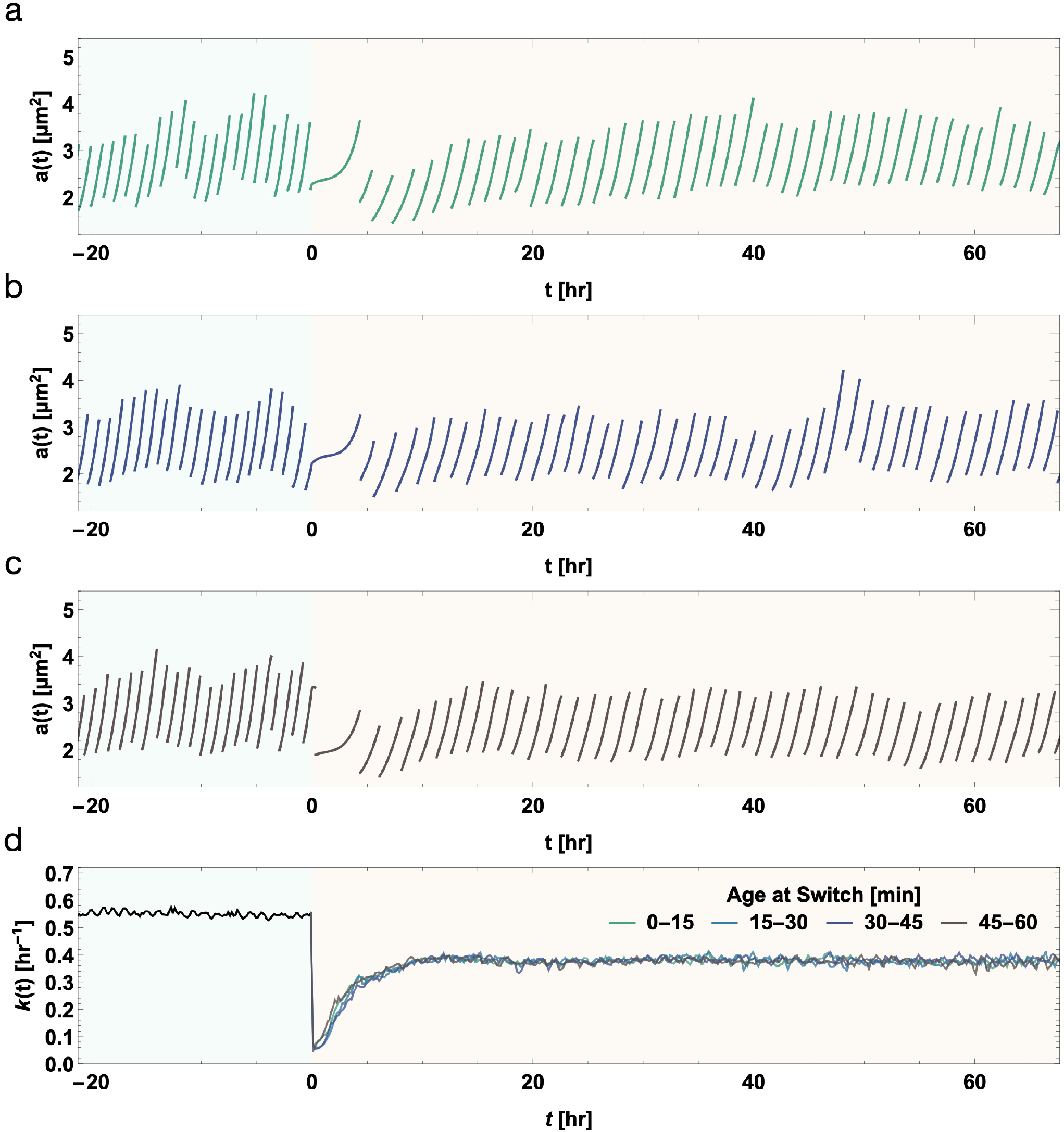
Data collapse of transient instantaneous growth rates of different cells experiencing abrupt shift in conditions at different ages. Cells growing in complex media (marked by blue background) in steady state were subjected to an instantaneous switch in growth media to simple media (marked by yellow background). **(a–c)** Typical area trajectory of a cell subjected to an instantaneous switch in growth conditions is shown through area versus time trajectories of actual individual cells growing and dividing over successive generations. Each cell trajectory encounters the switch in growth condition at a different age, and is colored accordingly (see (d)). **(d)** Cells were divided into four bins (marked with different colors) depending on their age at switch (the time between switch and the last division before it). The post-switch average instantaneous growth rate for each bin is plotted as a function of time. The overlap between the instantaneous growth rates from different bins indicates that the observed transient behavior of instantaneous growth rate post switch is unaffected by the age of the cells at switch.

**Fig. 7.**
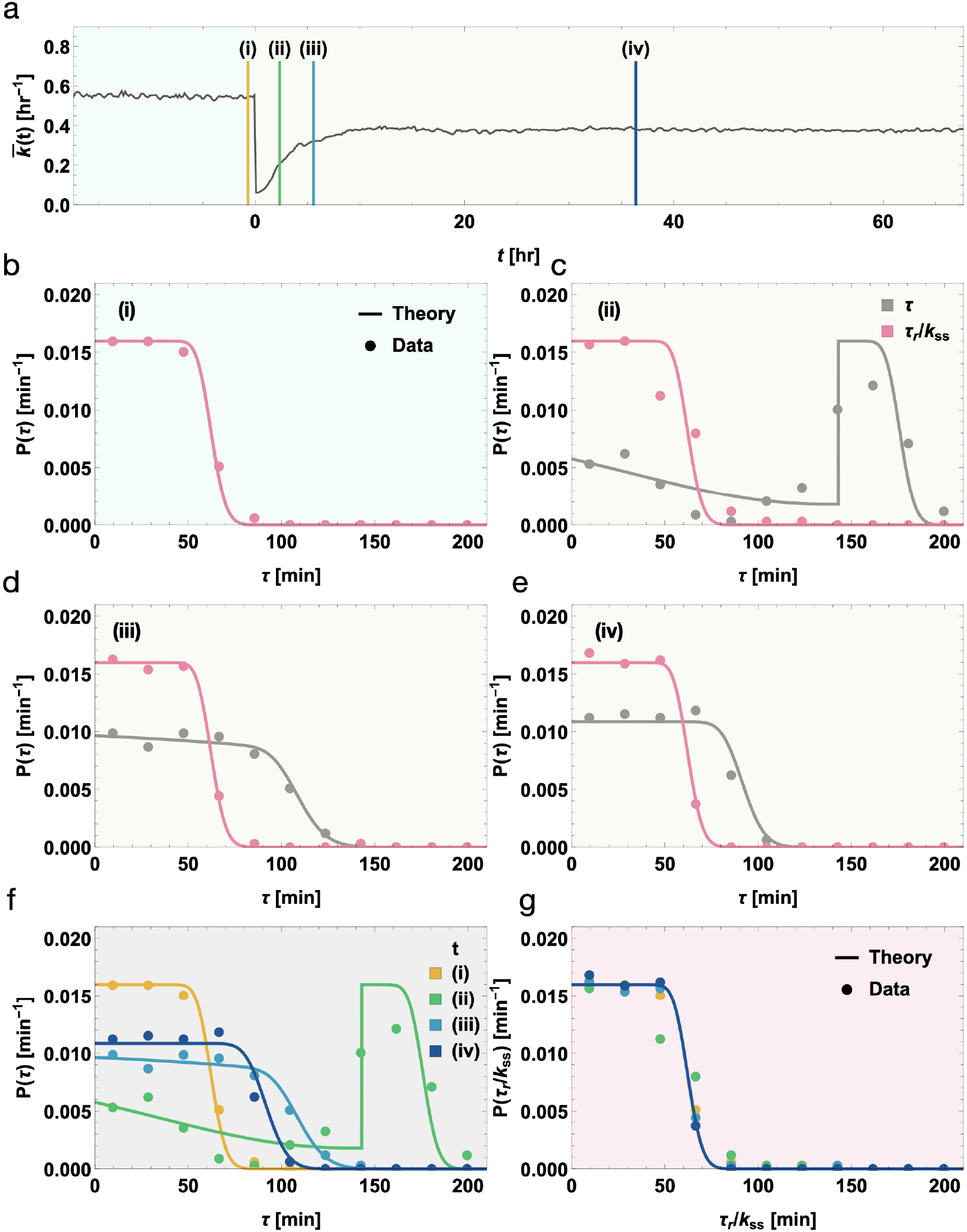
Dynamical rescaling by the instantaneous growth rate renders the dynamics time-invariant in the cellular frame of reference, as validated by the fitting-parameter-free data-theory matches of the cell age distributions. **(a)** The mean of the asynchronous instantaneous growth rate is plotted across the abrupt switch in growth conditions. This mean value is used along with the steady state division time distribution to predict the transient age distributions without any free parameters. **(b–e)** Distributions of cell ages at different experiment time instants corresponding to (i–iv) marked in (a) with colored lines. The points represent the experimentally measured distributions, while the solid lines are the theoretical predictions (see Methods section for details). The age distributions in both the lab frame of reference (*τ*, colored pink), and the cell’s intrinsic frame of reference (*τ*_*r*_, colored blue) are plotted. The distributions in the cell’s intrinsic frame of reference are scaled by a constant value (*k*_*ss*_, or the average steady-state growth rate before switch) to aid in visual comparison. **(f–g)** The age distributions at the different time points are shown together, in (f) the lab frame of reference, and (g) the cell’s intrinsic frame of reference. Theory predicts that the age distribution in the cell’s intrinsic frame of reference does not vary with time, which also matches the experimental observation, despite the large variations in the age distribution in the lab frame of reference during the transient period.

**Fig. 8.**
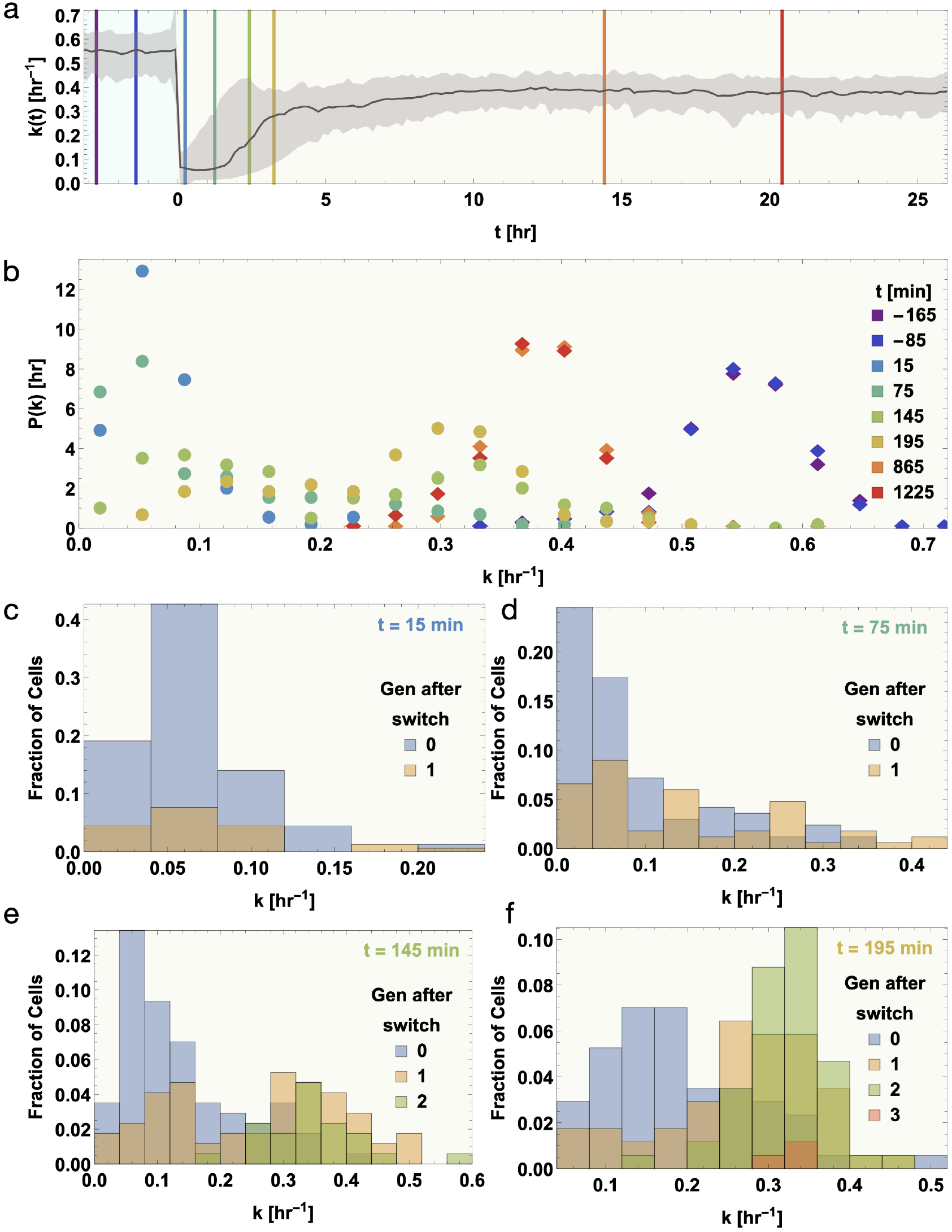
Transient dynamics following a change in environmental conditions yield bimodal transient single-cell growth-rate distributions. **(a)** The median instanta-neous growth rate is plotted as a function of time. The gray region around the median shows the central 90% data range. **(b)** The instantaneous growth-rate distributions are plotted at different times after switch, corresponding to the time points marked in (a) by lines of corresponding colors. The initial and final steady-state distributions are marked by solid rhombi, while the transient distributions are marked by circles. The final steady-state distributions have a larger coefficient of variation than the initial ones. We see bimodality in the transient distributions, and as time progresses, the height of the mode at smaller *k* decreases along with a corresponding increase in the height of the other mode at larger *k*. **(c–f)** The cause of the bimodality is investigated by plotting the fraction of cells with different growth rates at each time point, separated according to the number of divisions the cell underwent since the instantaneous switch in growth media. The zeroth generation corresponds to the cell cycle during which the switch happened, and each subsequent division increases the generation number by 1. It is clear that cells in their zeroth generation all belong to the mode with lower *k*, while the mode with greater *k* consists of cells that have undergone one or more divisions. Cells with multiple divisions almost exclusively belong to the mode with the greater *k*.

### Preliminaries and notation

In (14, 23) we identified the *intra*generational scaling laws or emergent simplicities governing stochastic growth and division of individual bacterial cells in *constant* growth conditions. We showed that a growth condition-specific emergent cellular unit of time governs stochastic growth and division. This timescale simply assumes a different numeric value from condition to condition while the rescaled (dimensionless) dynamics remain universal. Consistently, the interdivision-time distribution, *P* (*τ*), when mean-rescaled, is universal across conditions (14). Here *τ* is interpreted as the inter-division time interval. Bridging the scales of single-cell and population growth dynamics, we have also shown the equivalent result that the cell-age distribution, *G*(*τ*), when mean-rescaled, is universal for an asynchronous population in balanced growth conditions (16). Here, cell age *τ* is interpreted as the time elapsed (“age”) since the last division event. In constant growth conditions, the pre-divisional cell size increases exponentially on average (14); we denote this generational individual-cell (size) growth rate by *k*. Since *C. crescentus* cells divide asymmetrically, the single-cell (size) growth rate and the population (number) growth rate cannot be used interchangeably; however, we can write down the precise relation connecting them (16, 23, 24). Going beyond constant growth conditions, when environmental conditions vary with time, cell sizes may increase non-exponentially during inter-division intervals (for instance, as shown in Figs 6); we will use the instantaneous growth rate, *k*(*t*), defined as the time derivate of the logarithm of the cell size, to characterize transient growth dynamics in time-varying conditions.

## Results and discussions

### (I) The single-cell size growth rate is in stochastic intergenerational homeostasis in constant growth conditions

In balanced growth conditions, the individual-cell size growth rate is effectively a generational constant (14, 23). However, its value fluctuates from generation to generation even under constant growth conditions. How then can one establish whether it is in stochastic intergenerational homeostasis? While constancy of the generational growth rate’s mean value (averaged over the asynchronous population of cells in the single-cell experiment at each instant of time) is necessary (Fig 2(a)), this condition alone does not suffice; the mythical average cell does not capture single cell dynamics (25). The sufficient condition is that the *distribution* of generational single-cell growth rates sampled from different time windows of the experiment must remain invariant (17, 26). As shown in Fig. 2, we directly establish that this condition is met. Hence we conclude that the generational single-cell growth rate maintains stochastic intergenerational homeostasis under constant growth conditions.

### (II) The intergenerational dynamics of the individual– cell (size) growth rate are non-Markovian

Does the individual cell’s size growth rate serve as a repository of intergenerational memory? To answer this question unequivocally, we must disentangle the signatures of memory retention from the inherent stochasticity of the intergenerational process (27, 28). As is well known, when a process is memory-free (history-independent) or Markovian, future excursions of the system are determined solely by the present value of the stochastic variable, irrespective of history (27). Conversely, if the intergenerational temporal evolution of the single-cell growth rate in constant growth conditions is memoryful or non-Markovian, the history dependence can be revealed by considering the families of conditional probability distributions: *P* (*k*_*n*+1_ | *k*_*n*_, *k*_*n−*1_, …), where *k*_*n*_ denotes the value of the growth rate in generation *n* and *P* (*k*_*n*+1_ | *k*_*n*_, *k*_*n−*1_, …) denotes the probability that the growth rate in the (*n* + 1)^th^ generation is *k*_*n*+1_, given that the rates in the previous generations have values (*k*_*n*_, *k*_*n−*1_, …). For tractability, we consider the sequence of conditional probabilities *P* (*k*_*n*+1_ | *k*_*n*_, *k*_0_), whose *k*_0_-dependence characterizes non-Markovianity due to the *n*^th^ past generation. These are shown in Fig. 3 for generational indices *n* = 1 to *n* = 42, where *k*_*n*_ and *k*_0_ have been discretized to binary variables that are either above or below the median, 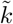,of the steady state distribution. By inspection, the differ-ence between 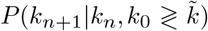, namely, the difference between the solid and dashed curves of the same color in Fig. 3, persists for tens of generations. Thus it is clear that *the cell growth rate obeys non-Markovian dynamics with a history dependence extending over tens of generations, even under constant growth conditions*. This behavior contrasts sharply with the Markovian memory-free or elastic character of the intergenerational homeostasis of individual bacterial cell sizes (17, 26).

### (III) Quantification of degree of non-Markovianity of intergenerational single-cell growth-rate dynamics

To characterize the degree of non-Markovianity of the single-cell growth rate, we introduce and define the following measure:

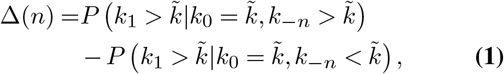

where, 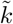 is the median single cell growth rate, and *k*_*i*_’s are the cell cycle growth rates of the *i*^*th*^ generation. For a completely Markovian intergenerational process, the probability distribution of *k*_1_ would depend only on *k*_0_ and be independent of *k*_−*n*_, thus, Δ(*n*) would be zero for all positive integers *n*. Thus, Δ(*n*) is a quantification of the non-Markovian influence of the growth rate *n* + 1 generations prior on the current generation’s growth rate, after excluding the Markovian influence of the previous generation’s growth rate. Fig. 4 shows that Δ takes around 40 generations to go to zero, indicating that single cell growth rate retains memory for ∼40 generations. The positive values of Δ for *n <* 40 indicate a positive correlation of growth rate in the current generation with growth rates of past generations, after compensating for the effect of growth rate in the previous generation.

### (IV) Distinct signatures of long- and short-term memory retention in the persistence curve

As shown in Figs. 3 and 4, the magnitude of the non-Markovian effect gradually decreases over tens of generations. To characterize memory persistence (27–29), or, equivalently, the “forgetting” of the past, we plot the generation–generation autocorrelation function of individual cell growth rates in constant growth conditions in Fig. 5. We show the results using linear and log-linear plots, and log-log plots for multiple growth conditions. The long range and slow decay of the autocorrelation function, extending to tens of generations, is evident for all the growth conditions (Fig. 5 d–f). The autocorrelation function displays a clear biphasic behavior, which we interpret as signatures of short- and long-term memory retention, suggesting that distinct underlying biological mechanisms are responsible for short-term (working) memory and long-term (fossilized) memory. There is a prominent “knee” separating the two regimes at *n* ∼ 20 generations. Two putative power-law regimes are revealed in the log-log plots, with the caveat that caution must be exercised in interpreting biological data as showing power-law behaviors (30, 31). The power law exponents are comparable for all conditions, and are tabulated in the Supplementary Information.

### (V) The instantaneous specific growth rate encapsulates universal signatures of plastic adaptation of individual cells subjected to time-varying environmental conditions

Now we turn our attention to *dynamic* growth conditions. We focus on the experimental scenario in which an asynchronous population of cells is initially in steady state in a constant growth condition (complex nutrient media). These cells subsequently undergo an abrupt shift to a novel condition (minimal nutrient media), which they have never previously encountered in their (multigenerational) lifetimes. We observe these cells for tens of generations after the abrupt change in conditions—long enough for each cell to attain a new steady-state. Since we started out with an asynchronous population, different cells experience the switch in conditions at different ages. At first glance, the results display bewildering complexity as different cells follow different non-exponential growth curves during the transient dynamics immediately following the change in conditions (Fig. 6 a–c). Surprisingly then, upon extracting the corresponding instantaneous growth rates (the time derivative of the logarithm of the cell size), the data collapse onto a single curve (Fig. 6 d)—an emergent simplicity! Thus the challenge in extracting dimension reduction from these complex and stochastic dynamics is conveniently overcome (32–35). Furthermore, the fact that the instantaneous growth rate serves as a direct and universal calibration of how the cell population experiences dynamic changes in their environment has the following remarkable implication. While the cell-age distributions sampled at different times since the switch differ dramatically in shape as they adjust from the previous steady-state to the new conditions, in the appropriately rescaled “cellular frame of reference” the correspondingly rescaled cell-age distributions remain time-invariant, undergoing a spectacular scaling collapse! The import of this observation is that the previously known scaling law for mean-rescaled age distributions in balanced growth conditions (see Preliminaries section) is a special instantiation of a much more general result: a dynamic rescaling of the cellular unit of time by the instantaneous growth rate captures the predominant effect of external variations in conditions (14, 16, 36). Using this temporal scaling result, we can derive exact analytic results for how the time-dependent cell-age distribution adapts to changing conditions (36). As shown in Fig. 7 the theory–data matches are compelling, despite the absence of any fine-tuning fitting parameter. *Our results thus reveal the natural representation for these time-dependent dynamics*.

### (VI) Bimodal distribution of instantaneous specific growth rates of individual cells during stochastic and plastic adaptation to time-varying environments

No-tably, as shown in Fig. 8 a–b, the distribution of instant-neous growth rates transitions from the initial to the final steady state forms, which are tight unimodal distributions, through intermediate *transient bimodal distributions*. Additionally, the final steady state distribution has a larger coefficient of variation (CoV) than the initial (see Fig. 11) indicating sloppier control over growth-rate homeostasis while in the minimal media after switch. Immediately following the change in conditions, the growth-rate distribution becomes bimodal during the transition, with a strong mode at lower values immediately after the switch. Thereafter, locations of the modes remain approximately constant, but the weight gradually shifts from the lower to the higher mode, suggesting a binary rather than a graded response. Further investigations into the process by which this transition occurs, accom-plished by decomposing the total distribution into subpopulations determined by generation after switch, are shown in Figs. 8 c–f and 9 a–b. We find that the growth-rate distribution is strongly generation-dependent immediately after the switch. At the same instant, all cells that have not divided since the switch have growth rates in the lower mode, cells two generations after the switch all correspond to the higher mode, and cells one generation after the switch are split between the two modes (Fig. 8 c–f). Some of this effect can be attributed to the strong memory in growth rate. Since cells with higher growth rate divide faster, a significant portion of cells with larger generation numbers post-switch are those that had higher growth rate initially, and retained this higher growth rate later (Fig. 9 a–b). But should this be the only factor underlying this effect, it would fail to explain the shift in weight between the modes—this occurs due to the resetting of the growth rate value to the higher mode. Indeed, we find another contributing factor: during the transient recovery period, among the cells starting with similar growth rates at the initial time, those that divide are more likely to have higher growth rates at a later time, indicating a finite jump-induced recovery of growth rate after division (Fig. 9 c). This finite post-division jump causes the shift in weight between the modes. Within all generations during the transient period of growth rate adjustment, growth rates gradually increase with time, though the final steady-state growth rate is distributed over lower values than the initial steady-state growth-rate distribution.

**Fig. 9.**
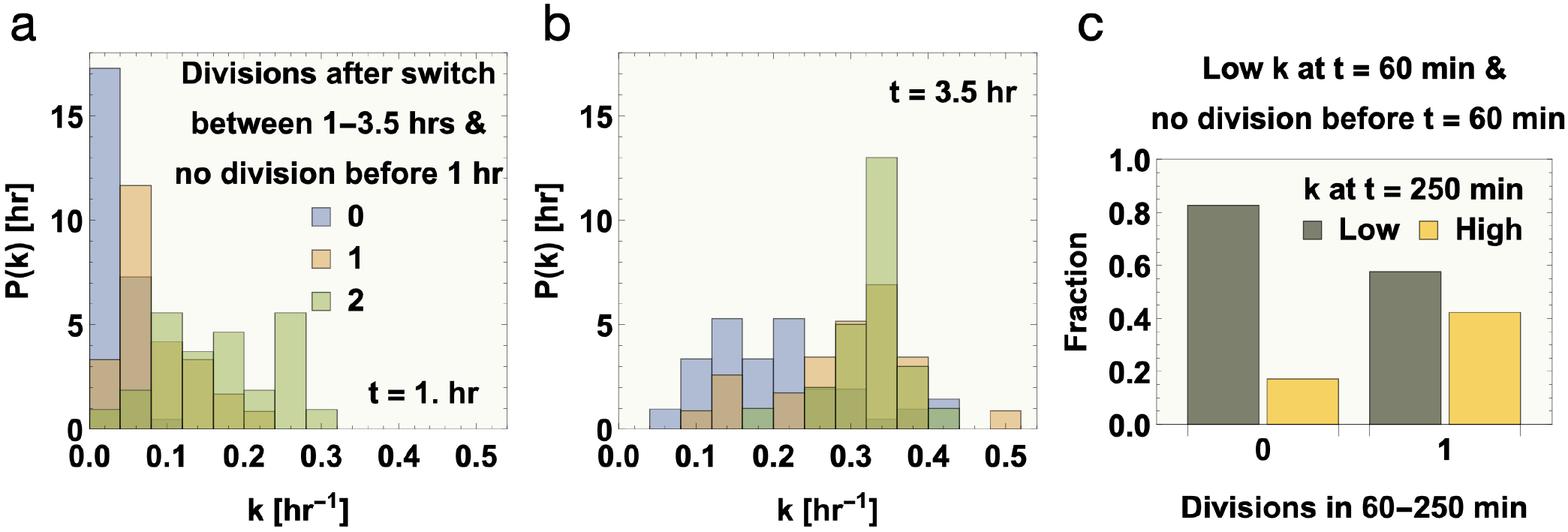
Establishing the link between memory in growth rate, number of cell divisions in a given time window, and the growth rate at the end of the time window, during the recovery period after an instantaneous switch in growth conditions. **(a-b)** The instantaneous growth rate (*k*) distributions of cells divided into 3 bins according to the number of divisions in the time window of 1–3.5h are plotted at (a) the start of and (b) the end of the time window. Only those cells which did not undergo any division before 1 h after the switch are considered. These distributions indicate that cells that had more divisions in a given time interval generally have a higher *k* both at the start and at the end of the time interval. Thus, cells retain a strong memory of *k*, and cells with higher *k* are more likely to divide. **(c)** Here, we investigate the effect of division on the recovery of *k* after the switch, after excluding the effect of memory retention in *k* by only considering cells that start out with low *k* (i.e., cells with *k* below the median) at the start of the time interval of 60–250 min after the switch. The fraction of these cells that stay in the low *k* range or switch to the high *k* range at the end of the interval are plotted for cells that underwent no division in the interval, and those that underwent one division. The fraction of cells switching from low to high *k* are much higher for cells that underwent one division, indicating that division helps in the recovery of *k* after the switch. The reason for choosing a longer interval in (c) compared to (a–b) is to ensure the availability of a sufficient number of data points for the selected conditions considering that cells that start out with low *k* take longer to divide.

**Fig. 10.**
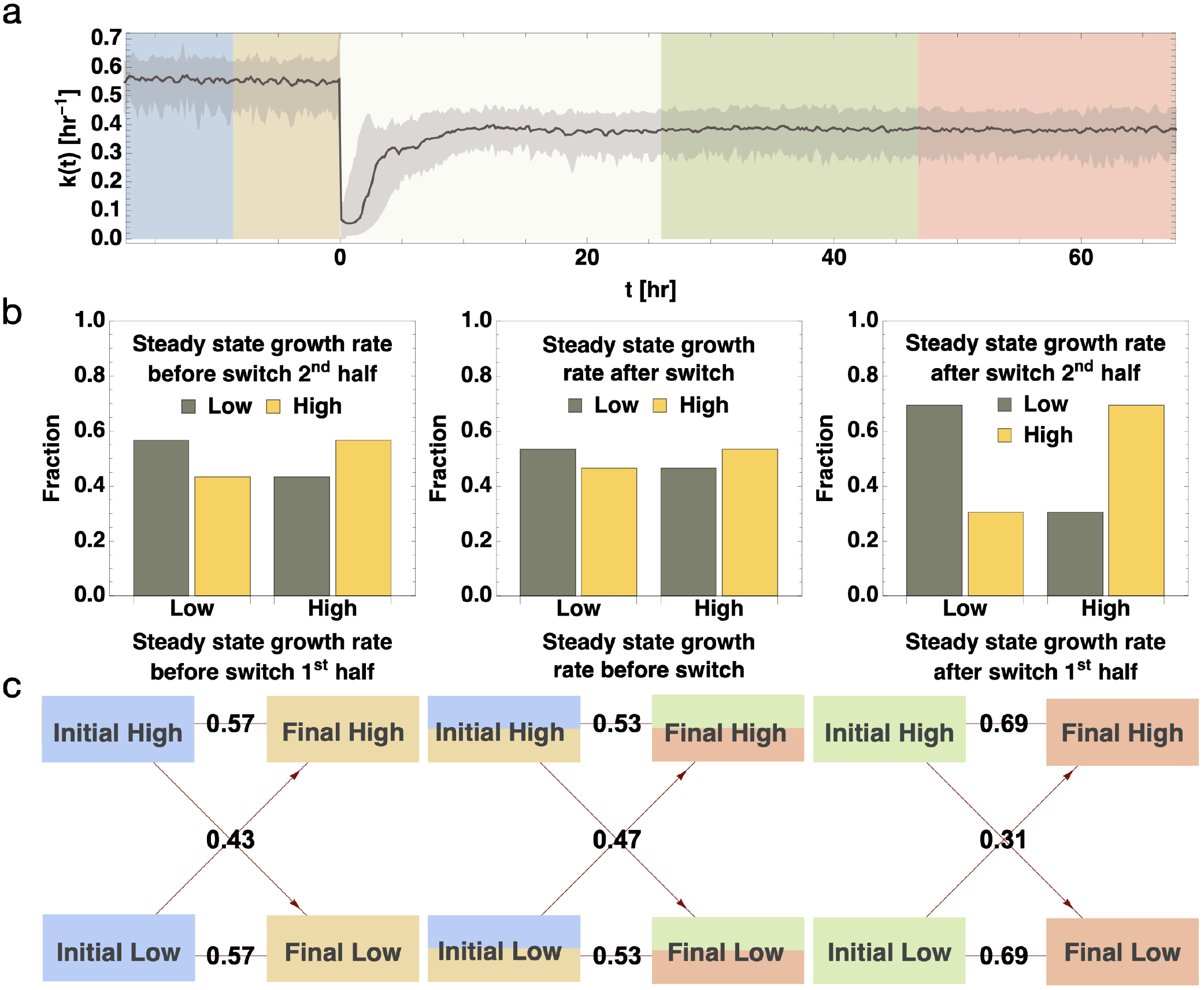
Interplay between retention and resetting of a cell’s memory of its growth rate in dynamic conditions. **(a)** The median instantaneous growth rate is plotted as a function of time for cells subjected to an abrupt shift in nutrient condition at *t* = 0, along with colored regions depicting the time intervals when growth rate is in steady state. Blue and yellow correspond to the first and second halves of the steady-state region before the switch, and green and red to the first and second halves of the steady state region after the switch. **(b–d)** Bar charts showing the probability for a cell with low (below median) or high (above median) growth rate in one region to transition to low or high growth rate in another region. The directed graphs below the bar charts explicitly display the corresponding transition probabilities. These are plotted for transitioning between (a) the two halves of the steady state before switch (blue to yellow), (b) steady state before switch to steady state after switch (blue and yellow to green and red), and (c) the two halves of the steady state after switch (green to red). The larger the difference from 0.5 in the transition probabilities, the greater the memory retention. Thus, compared to within the steady state periods before and after the switch, long-term memory is reset across the switch.

**Fig. 11.**
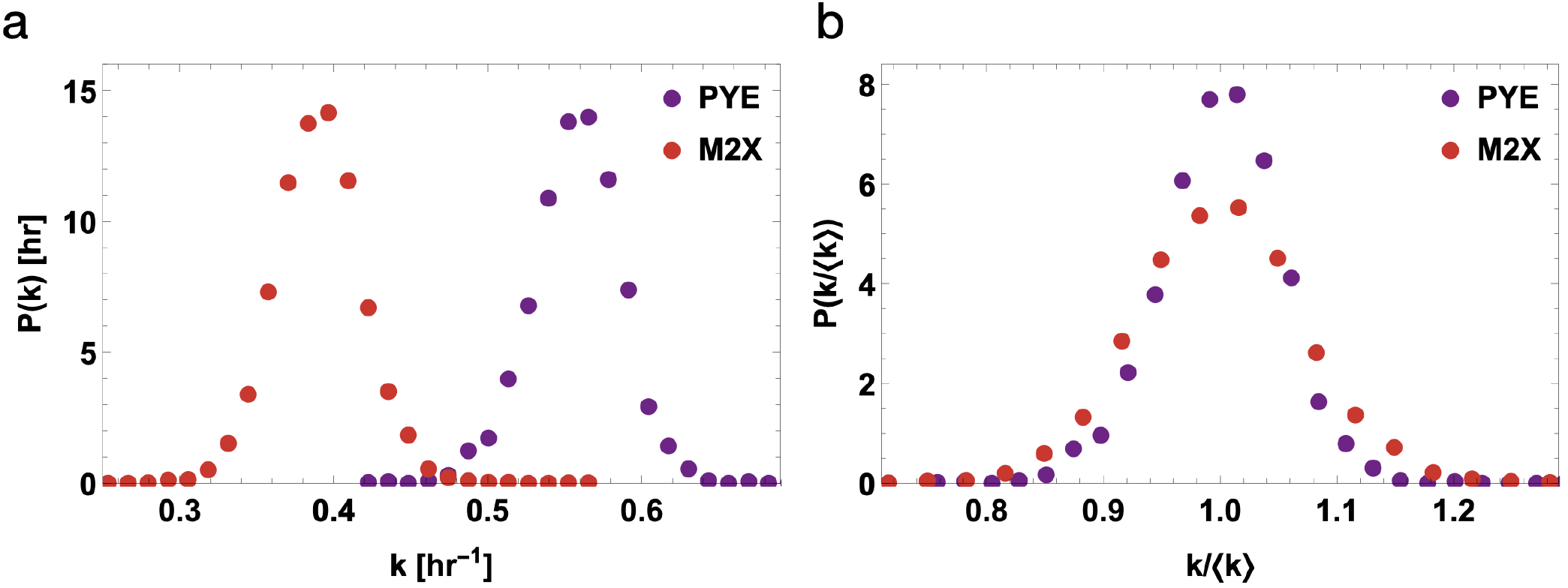
Increased variability in growth-rates after a new homeostasis is achieved following a change in environmental conditions. **(a)** The growth-rate distributions are plotted for the steady-state periods before and after the instantaneous switch from complex (PYE) to minimal (M2X) media. The steady-state time periods are the same as those shown in Fig. 10. **(b)** The distributions in (a) rescaled by their corresponding mean values are shown. The rescaled distribution is wider for the after-switch minimal media (M2X), indicating an increase in CoV.

### (VII) Retention and resetting of a cell’s memory in constant and variable environments

As noted previously, we find that even under constant environmental conditions, non-Markovian memory of the stochastic intergenerational dyamics of the individual-cell growth rate is retained over tens of generations; the persistence curve extends over a similar intergenerational timescale in constant growth conditions. The question naturally arises as to how the non-Markovianity is expressed and reset in dynamical growth conditions. For a more long-term intergenerational estimate of the non-Marovian memoryful evolution of the growth rate, in Fig. 10 we characterize the slow mixing of the growth rate across the environmental shift. This is accomplished by dividing the cell population into two halves at every instant based on how the individual growth rate relates to the median of the corresponding steady-state growth-rate distribution, and characterizing the amount of non-ergodic mixing between any two time points by the amount by which the transition probabilities between the two halves of the population differ from 0.5. This measure directly reflects the extent of correlation between growth rates at those two time points. From the analysis shown in Fig. 10, it is clear that the transition across the switch is more ergodic than within each of the steady states on both sides of the switch. In conjunction with Fig. 9 and the discussion in the last section, this is strong evidence of a *resetting* of long-term memory across the switch.

## Concluding remarks

We have previously delineated two distinct schemes by which stochastic homeostasis may be achieved and sustained in self-regulating control systems: passive “elastic adaptation”, achieved through reflexive memory-free responses to instantaneous conditions (as exemplified by Ashby’s Homeostat, the first technical model of homeostasis ever realized) or active “plastic adaptation”, achieved through sophisticated control based on past experiences to achieve reflective responses (as exemplified by Shannon’s “Maze Solving Machine” which stored memories of trajectories from past excursions) (17, 37, 38). In (17, 26) we have shown that growing and dividing individual bacterial cells maintain stochastic intergenerational cell-size homeostasis in constant growth conditions through elastic adaptation, i.e., Markovian or memory-free dynamics. In contrast, in this work, we have shown that the intergenerational cell growth-rate dynamics are non-Markovian, or “plastic” and memoryful, under steady-state conditions. Using the metric of non-Markovianity introduced in Eq. Eq. (1), we have definitively established that strong non-Markovian memory is retained for tens of generations. The generational autocorrelation function of the single-cell growth rate, the persistence curve, also validates that intergenerational memory of the instantaneous growth rate persists for tens of generations in constant growth conditions. The two regimes observed in the persistence curve suggest a delineation between short-term(working) and long-term (fossilized) memories in the cellular growth rate, characterized by differing characteristics of memory loss or “forgetting”. Since the single-cell growth rate retains longterm memory of the nutritional history, its instantaneous or generational values do not specify cellular phenotypes (39–41).

Despite the complex non-Markovian factors involved in determining the instantaneous growth rate at any instant in time, we find the surprising emergent simplicity that once the growth rate has been determined as a function of time, the complex cellular dynamics of growth and division under time-varying conditions can simply be gleaned from it through an appropriate rescaling of time. In the intrinsic cellular frame of reference thus obtained, the growth and division dynamics always remain in steady state despite any transient changes taking place in the lab frame of reference. Under dynamic growth conditions characterized by an abrupt shift in conditions, during the transient period as the cell adapts to a novel condition, it “resets” its long-term memory of growth rate. This remarkable result suggests that “flushing” of long-term memory plays an important role in physiological adaptation (29, 42–44). Our results challenge the notion that the primary goal of physiological regulation in sustaining homeostasis is to achieve error correction through noise suppression mechanisms, thus holding key physiological variables at desirable set-point values (9). In fact our results for transient adjustments to the instantaneous growth-rate in time-varying conditions explicitly show that the initially narrowly-distributed growth rates become noisy bi-modal distributions during transient adjustments to dynamic conditions, and ultimately achieve a new homeostasis with a broader distribution, i.e., increased variability.

It is increasingly appreciated that unicellular organisms can serve as exciting arenas for explicating principles governing statistical learning in neuron-free naturally occurring complex systems (45–48). In this work we have established that plastic adaptation of an individual cell’s instantaneous growth rate involves non-Markovian memory and history-dependent physiological regulation, resulting in increased variability (broader growth rate distribution) even after the transients have died out and a new homeostatic steady state is reached. Whether this increased variability leads to improved predictions and anticipation through cellular learning mechanisms attuned to negotiating repeating patterns in dynamic changes in environment is an outstanding open question.

## Methods

### Experimental methods

Cells of the conditionally adhesive *C. crescentus* strain IB001 growing in complex medium (peptone-yeast extract; PYE) were introduced into a single-channel microfluidic device and incubated in the presence of the vanillate inducer (which promotes adhesion) for one hour. Then, complex growth medium was infused through the channel at a constant flow rate of 10 *μ*L/min (Pump 11 Elite, Harvard Apparatus) to flush out non-adherent cells and swarmer daughter cells born throughout the course of the experiment, thereby preventing over-crowding. This also ensured continuous delivery of nutrients and removal of waste.

A microscope (Nikon Ti Eclipse with perfect focus system) and motorized XY stage (Prior Scientific ProScan III) under computerized control (LabVIEW 8.6, National Instrument) were used to acquire phase-contrast images at a magnification of 250X (EMCCD: Andor iXon+; objective: Nikon Plan Fluor 100X oil objective plus 2.5X expander; lamp: Prior LED). The microfluidic device, growth media, and imaging apparatus were located inside a custom-built acrylic microscope enclosure equilibrated to 30.0^°^C with temperature controller; this chamber was in turn located within a sealed environment with defined temperature and humidity for an additional level of environment control. Image analysis to extract single-cell growth trajectories was conducted using custom-built Python software.

For the experiment in which cells were subjected to an instantaneous change in growth condition, after 24 hours of infusion of complex medium, the growth medium was instantaneously changed to minimal medium with xylose as the sole carbon source (M2X). This change was executed automatically by pre-programming the pump delivering complex medium through one of the inputs to the microfluidic channel to stop after 24 hours, and another pump to simultaneously start pumping minimal medium through another input at the same flow rate.

### Theoretical methods

For a cell of a given age (*τ*) at any given lab time (*t*), its age in the intrinsic cellular frame of reference is given by,

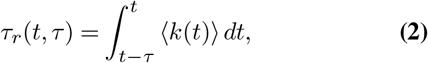

where ⟨*k*(*t*)⟩ is the average instantaneous growth rate experimentally measured as a function of time (Fig. 7 a). The theoretical age distribution in the intrinsic cellular frame of reference, after being scaled by 1*/k*_*ss*_ (where *k*_*ss*_ is the average steady state growth rate from before the switch in growth conditions), is given by,

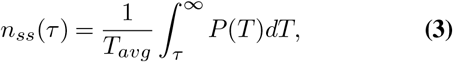

where *P* is the experimentally measured steady state division time distribution before switch, and *T*_*avg*_ is its mean. The theoretical age (*τ*) distribution in the lab frame of reference at lab time *t* is given by,

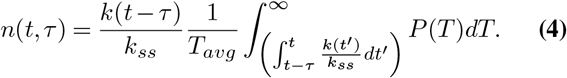

The derivations of these equations proceed as described in (36).

## ACKNOWLEDGMENTS

We thank Purdue University Startup funds, Purdue Research Foundation, the Purdue College of Science Dean’s Special Fund, and the Showalter Trust for financial support. K.J., K.F.Z., and S.I.-B. acknowledge support from the Ross-Lynn Fellowship award. R.G., J.S., and S.I.-B. thank the National Science Foundation (NSF REU Grant No. PHY-1460899) for financial support. We are grateful to Iyer-Biswas group members for insightful discussions, assistance with image analysis, and feed-back on the manuscript. The datasets for constant growth conditions at 24^*°*^C and 34^*°*^C are published in (14).

## AUTHOR CONTRIBUTIONS

C.S.W, R.R.B., and S.I.-B. conceived of the research; K.J., K.F.Z., C.S.W, R.R.B., and S.I.-B. designed the research; K.J. spearheaded data analysis and theory development under the guidance of S.I.-B.; K.J., S.R., R.R.B. and S.I.-B. performed analytic calculations; K.F.Z refined and redesigned the experimental setup; K.F.Z, C.S.W. and S.I.-B. performed experiments; S.R. performed image analyses with inputs from K.J., C.S.W. and S.I.-B.; R.G. and J.S. contributed to image analysis under the guidance of S.R. and S.I.-B.; K.J., C.S.W., R.R.B., and S.I.-B. wrote the paper; S.I.-B. supervised the research.

